# The demyelinating agent cuprizone induces a male-specific reduction in binge eating in the binge-prone C57BL/6NJ strain

**DOI:** 10.1101/865600

**Authors:** Richard K. Babbs, Jacob A. Beierle, Julia C. Kelliher, Rose Medeiros, Jeya Anandakumar, Anyaa Shah, Emily J. Yao, Melanie M. Chen, Camron D. Bryant

## Abstract

Binge eating (**BE**) is a heritable symptom of eating disorders with an unknown genetic etiology. Rodent models for BE of palatable food permit the study of genetic and biological mechanisms. We previously used genetic mapping and transcriptome analysis to map a coding mutation in *Cyfip2* associated with increased BE in the BE-prone C57BL/6NJ substrain compared to the BE-resistant C57BL/6J substrain. The increase in BE in C57BL/6NJ mice was associated with a decrease in transcription of genes enriched for myelination in the striatum. Here, we tested the hypothesis that decreasing myelin levels with the demyelinating agent cuprizone would enhance BE. Mice were treated with a 0.3% cuprizone home cage diet for two weeks. Following a three-week recovery period, mice were trained for BE in an intermittent, limited access procedure. Cuprizone induced similar weight loss in both substrains and sexes that recovered within 48 h after removal of the cuprizone diet. Surprisingly, cuprizone reduced BE in male but not female C57BL/6NJ mice while having no effect in C57BL/6J mice. Cuprizone also reduced myelin basic protein (**MBP**) at seven weeks post-cuprizone removal while having no effect on myelin-associated glycoprotein (**MAG**) at this time point. C57BL/6N mice also showed less MBP than C57BL/6J mice. There were no statistical interactions of Treatment with Sex on MBP levels, indicating that differences in MBP are unlikely to account for sex differences in BE. To summarize, cuprizone induced an unexpected, male-specific reduction in BE which could indicate sex-specific biological mechanisms that depend on genetic background.

## INTRODUCTION

Binge eating (**BE**) is operationally defined as the consumption of a relatively large amount of food over a short period of time, and is accompanied by feelings of loss of control (Amianto, Ottone, Abbate Daga, & Fassino, 2015). Repeated BE that occurs at least once per week for at least three months is termed binge eating disorder (**BED**). BE has a lifetime prevalence of approximately 4.5% (Hudson, Hiripi, Pope, & Kessler, 2007) and is associated with a number of comorbid behavioral, mood, and physical disorders such as substance abuse, depression, obesity, and chronic pain (Bulik & Reichborn-Kjennerud, 2003; Citrome, 2017; Hudson et al., 2007). Although there is no sex difference in the lifetime prevalence of BE, women are nearly twice as likely as men to progress to the more severe BED (3.5% vs. 2.0% respectively (Hudson et al., 2007). However, the etiology of this sex difference is not known.

Numerous neurophysiological and neuroanatomical changes have been identified in eating disorders (ED), including BED, bulimia nervosa (**BN**) and anorexia nervosa (**AN**), including alterations in both dopaminergic and serotonergic signaling in BN and AN (Frank et al., 2005; Kaye, 2008; Tauscher et al., 2001), reduced cerebral glucose metabolism in the anterior cingulate (Kojima et al., 2005; Naruo et al., 2001), and enhanced cerebral perfusion in the thalamus and amygdala-hippocampus complex in AN, and greater frontal cortical response to food stimuli in BED (Geliebter et al., 2006). Another important neuroanatomical change common to multiple types of ED involves changes in myelin which is the lipid-rich, membranous insulation sheath that extends from oligodendrocytes to the axons and is involved in axon maintenance (via oligodendrocyte signaling) and function, including rapid salutatory conduction of action potentials. BN is associated with a reduction in fractional anisotropy, a brain imaging measure of white matter integrity, in the corona radiata and corpus callosum (Mettler, Shott, Pryor, Yang, & Frank, 2013) as well as in forceps major and minor, inferior fronto-occipital, superior longitudinal, and uncinate fasciculi, corticospinal tract, and cingulum (He, Stefan, Terranova, Steinglass, & Marsh, 2016). In women with AN, a reduced fractional anisotropy relative to controls was reported in left superior longitudinal fasciculus (Via et al., 2014). Importantly, previous results found no differences in white matter integrity between healthy controls and women who have recovered from AN (Yau et al., 2013), indicating that decreased myelination is a reversible consequence rather than a cause of aberrant eating behavior in ED. Additionally, both obese women and men show a decrease in myelination in obesity, but only in women did reduced myelin correlate with BMI and leptin levels (Mueller et al., 2011), suggesting potential sex differences in the contribution of demyelination to negative health consequences of eating disorders.

Using transcriptome analysis via mRNA sequencing (**RNA-seq**), we previously showed that BE of sweetened palatable food (**PF**) relative to chow controls induced a downregulation of several genes enriched for myelination, axon ensheathment, and oligodendrocyte formation and differentiation, providing evidence that, similar to BN and AN, decreased myelination is a consequence of BE in our preclinical model (Kirkpatrick et al., 2017). However, because transcriptome analysis is conducted at the mRNA level, it is not known whether the BE-induced decrease in myelin-associated transcripts translates to a decrease in myelin proteins. Furthermore, whether decreasing myelin levels could induce or enhance BE has not been tested.

In the present study, we tested the hypothesis that administration of the demyelinating agent cuprizone (a.k.a. cyclohexylidene hydrazide) in C57BL/6 mice (Hiremath et al., 1998) would induce BE in the BE-resistant C57BL/6J (**B6J**) substrain and enhance BE in the BE-prone C57BL/6NJ (**B6NJ**) substrain (Kirkpatrick et al., 2017). Laboratory chow containing 0.3% cuprizone has previously been shown to induce a significant decrease in myelination of the corpus callosum at 4 weeks and complete demyelination after six weeks (Hiremath et al., 1998) and two-week exposure of 0.2% cuprizone has been shown to be sufficient to induce nearly complete demyelination in mice (Doan et al., 2013). At the completion of the study, we harvested brain tissue and quantified two myelin proteins that are downregulated in the mouse brain following cuprizone treatment, including myelin-associated glycoprotein (**MAG**) and myelin basic protein (**MBP**) (Ludwin & Sternberger, 1984).

## METHODS

### Mice

All experiments were conducted in accordance with the NIH Guidelines for the Use of Laboratory Animals and were approved by the Institutional Animal Care and Use Committee at Boston University (AN-15403). Mice were maintained on a 12 h /12 h light/dark cycle (lights on at 0630 h) and housed four animals per cage in same-sex cages. Laboratory chow (Teklad 18% Protein Diet, Envigo, Indianapolis, IN, USA) and tap water were available *ad libitum* in home cages prior to the experiment. Testing was conducted in the a.m. of the light phase (0800 h to 1200 h) on a 12 h light/dark cycle (lights on at 0630 h). 32 C57BL/6J (**B6J**) and 32 C57BL/6NJ (**B6NJ**) mice (7 weeks old) were purchased from the Jackson Laboratory (JAX; Bar Harbor, ME). Mice were habituated to the vivarium for one week prior to cuprizone treatment in a room separate from the testing room. Mice were 56 days old at the beginning of cuprizone treatment and 91 days old on the first day of BE training.

### Cuprizone diet

Sixty-four mice were assigned to a treatment group by cage (four mice per cage) in a 2 × 2 factorial design so that N = 15-16 (8 females, 8 males) per Substrain (B6J, B6NJ) per Treatment (0.3% Cuprizone-treated Chow, Untreated Chow). Untreated Chow mice continued to have *ad libitum* access to water and standard laboratory chow described above throughout the entire study. On D1 of the study, the remaining 32 mice had the standard chow replaced with a chow that was structurally and nutritionally identical, but also contained 0.3% cuprizone (Envigo, Indianapolis, IN, USA), a copper chelating agent that induces demyelination in C57BL/6 mice by causing mitochondrial and endoplasmic reticulum stress and apoptosis in mature oligodendrocytes that in turn generates myelin debris and activation of astrocytes and microglia (Gudi, Gingele, Skripuletz, & Stangel, 2014; Hiremath et al., 1998; Sen, Mahns, Coorssen, & Shortland, 2019). Mice were maintained on this diet for two weeks, and on D14 the cuprizone diet was replaced with standard laboratory chow as described above. Mice were allowed to recover for 21 days before BE training commenced. A nearly identical abbreviated, two-week cuprizone treatment protocol (0.2% rather than 0.3% cuprizone) was previously shown to be sufficient to induce robust demyelination in the corpus callosum at three weeks post-cuprizone treatment C57BL/6 (substrain not specified) mice (Doan et al., 2013). Specifically, the two-week 0.2% cuprizone regimen resulted in hallmark effects, including a marked depletion of mature oligodendrocytes and a significant accumulation of microglia and astrogliosis in the corpus callosum that peaked at the fifth week (3 weeks post-cuprizone termination) (Doan et al., 2013). The five week time point after 2 or 3 weeks of cuprizone diet is considered peak demyelination, after which remyelination begins and the levels of microglia and astrocytes decrease (Doan et al., 2013).

### BE procedure

Mice were trained in an intermittent, limited access procedure as described (R. K. Babbs et al., 2018; Richard K. Babbs et al., 2019; Kirkpatrick et al., 2017). We used a two-chambered conditioned place preference (**CPP**) apparatus, with differently textured floors in each chamber. The right and left sides were designated the food-paired and no-food-paired sides, respectively. Mice were trained and video recorded in unlit sound-attenuating chambers (MedAssociates, St. Albans, VT USA). On Day 1, initial side preference was determined by placing each mouse on the left, no-food-paired side with the divider containing an open entryway that provided free access to the both sides for 30 min. Clean, empty food bowls were placed in the far corners of each side. On Days 2, 4, 9, 11, 16, and 18, mice were confined to the food-paired side with a closed divider that prevented access to the no-food-paired side. Mice were provided forty, 20 mg sweetened palatable food pellets (**PF**; TestDiet 5-TUL, St. Louis, MO, USA) in a non-porous porcelain dish in the far corner of the chamber. On Days 3, 5, 10, 12, 17, and 19, mice were confined to the no-food-paired side with no access to the FP side. A clean, empty, and non-porous porcelain dish was placed in the far corner of the chamber during this time. On Day 22, side preference was again assessed with open access to both sides. No food was present in either bowl at this time. The experimenter was blinded to Genotype throughout data collection, video tracking, and analysis. Video-recorded data were tracked using AnyMaze (Stoelting Co., Wood Dale, IL USA).

### Light/dark conflict test of compulsive-like eating

Following BE training and assessment of PF-CPP, on Day 23, we subsequently assessed compulsive-like eating and concomitant behaviors in the anxiety-provoking light/dark conflict test (R. K. Babbs et al., 2018; Richard K. Babbs et al., 2019; Kirkpatrick et al., 2017) where rodents will normally avoid the aversive, light side. The light/dark box consists of a dark side, which is an enclosed black, opaque Plexiglas chamber, and a light side with clear Plexiglas. An open doorway allowed free access to both sides. A non-porous ceramic bowl containing forty, 20 mg PF pellets was placed in the center of the light side. Mice were initially placed on the light side facing both the food and the doorway and were video recorded for 30 min. Because the light side is aversive, increased behaviors in this environment were operationalized as compulsive-like, including compulsive-like eating.

On D24, brains were extracted and punches of the left and right striatum were harvested and combined as previously described (Kirkpatrick et al., 2017; Yazdani et al., 2015), flash frozen on acetone and dry ice, and stored at −80°C until western blotting procedures commenced. We chose to examine the striatum because this is the tissue in which we originally identified the downregulation of transcripts coding for genes associated with oligodendrocyte differentiation and myelination (Kirkpatrick et al., 2017). Cuprizone treatment has been shown to induce multiple signs of demyelination in the striatum, including the hallmark cuprizone effects of a decrease in MBP and other myelin proteins as well as an increase in astrogliosis (via GFAP) and microgliosis (via IBA1) in female C57BL/6N mice (Beckmann et al., 2018; Mandolesi et al., 2019)

### Immunoblotting for MAG and MBP

Striatal tissue was homogenized in RIPA buffer (Thermo Scientific, Waltham, MA USA, #89901) containing 1x HALT protease/phosphatase inhibitor cocktail (Thermo Scientific, #1861284) with an ultrasonic homogenizer and then spun at 17200 RCF for 20 min at 4°C to collect supernatant. Protein concentration was quantified by BCA protein estimation (Thermo Scientific, #23225). 30 μg of sample protein and loading buffer (BioRad, Hercules, CA #161-0747) was loaded into 4-15% Criterion TGX gels (BioRad, #5671085) and run at 200 V for 50 min. Gels were transferred onto nitrocellulose membranes (General Electric, Boston, MA, USA, #10600002) for 1 h at 90 V in Towbins buffer containing 20% methanol. Blots were then blocked with 5% BSA in TBST (0.5% Tween 20 in tris-buffered saline) for 1 h.

For MAG detection, blots were incubated in a 1:5,000 dilution of anti-MAG antibody (EMD Millipore, Burlington, MA, USA, #MAB1567) in 5% BSA in TBST overnight at 4C, then a 1:10,000 dilution of peroxidase conjugated donkey anti mouse antibody (Jackson Immunoresearch, West Grove, PA, USA, #715-035-151). For MBP detection, blots were incubated in a 1:10,000 dilution of anti-MBP antibody (EDM Millipore, #MAB 386) in 5% BSA in TBST overnight at 4C, then a 1:10,000 dilution of peroxidase conjugated donkey anti rat antibody (Jackson Immunoresearch #712-035-153) for one hour. After probing, all blots were imaged using Clarity ECL (BioRad, #170-5061) and a c300 imager (Dublin, CA, USA).

Blots were then stripped (Thermo Scientific, 46430) at 55°C for 30 min, reblocked with 5% BSA in TBST, incubated in 1:50,000 Beta-actin antibody (Sigma Aldrich, St, Louis, MO, USA, #A2228) in 5% BSA in TBST for 1 h, and then a 1:10,000 dilution of peroxidase conjugated donkey anti mouse antibody for 1 h. Blots were then reimaged for beta actin. Densitometry was conducted using ImageJ, and each lane was normalized to its respective densitometry value for beta actin. Each lane was then normalized to the average wild-type value of that particular blot and then finally, data were combined across blots and statistical analysis was run in R as described below.

### Statistical analysis

We analyzed food consumption in R (https://www.r-project.org/) using mixed model ANOVAs with Genotype, Sex, and Treatment as independent variables, and Day as a repeated measure. When the data were separated by Sex, mixed model ANOVAs were employed with Genotype and Treatment as independent variables, and Day as a repeated measure. Bonferroni-adjusted *post hoc* pairwise unpaired t-tests were used to determine whether group differences were significant on individual days. Slope analyses were conducted as previously described (R. K. Babbs, Wojnicki, & Corwin, 2012; Richard K. Babbs et al., 2019) using GraphPad Prism 7 (GraphPad Software, La Jolla, CA USA).

## RESULTS

### Cuprizone treatment induces weight loss in female and male B6J and B6NJ mice

We employed a regimen consisting of one week of habituation to the colony, two weeks of the cuprizone diet, three weeks of recovery from cuprizone, and four weeks of training for BE of PF (**Fig.1**). A single cuprizone-assigned male B6J mouse died early on during the study. Therefore, the results are presented for 63 mice instead of 64 mice. In examining the sex-combined dataset, we found significantly lower body weight in cuprizone-treated mice of both the B6J and B6NJ strains on D8-D14 of cuprizone treatment (**Fig.2A**). Remarkably, within 24 h after the cuprizone diet was replaced with standard laboratory chow (D15), mice regained nearly all of their body weight and did not differ significantly from control mice (**Fig.2A**).

**Figure 1.**
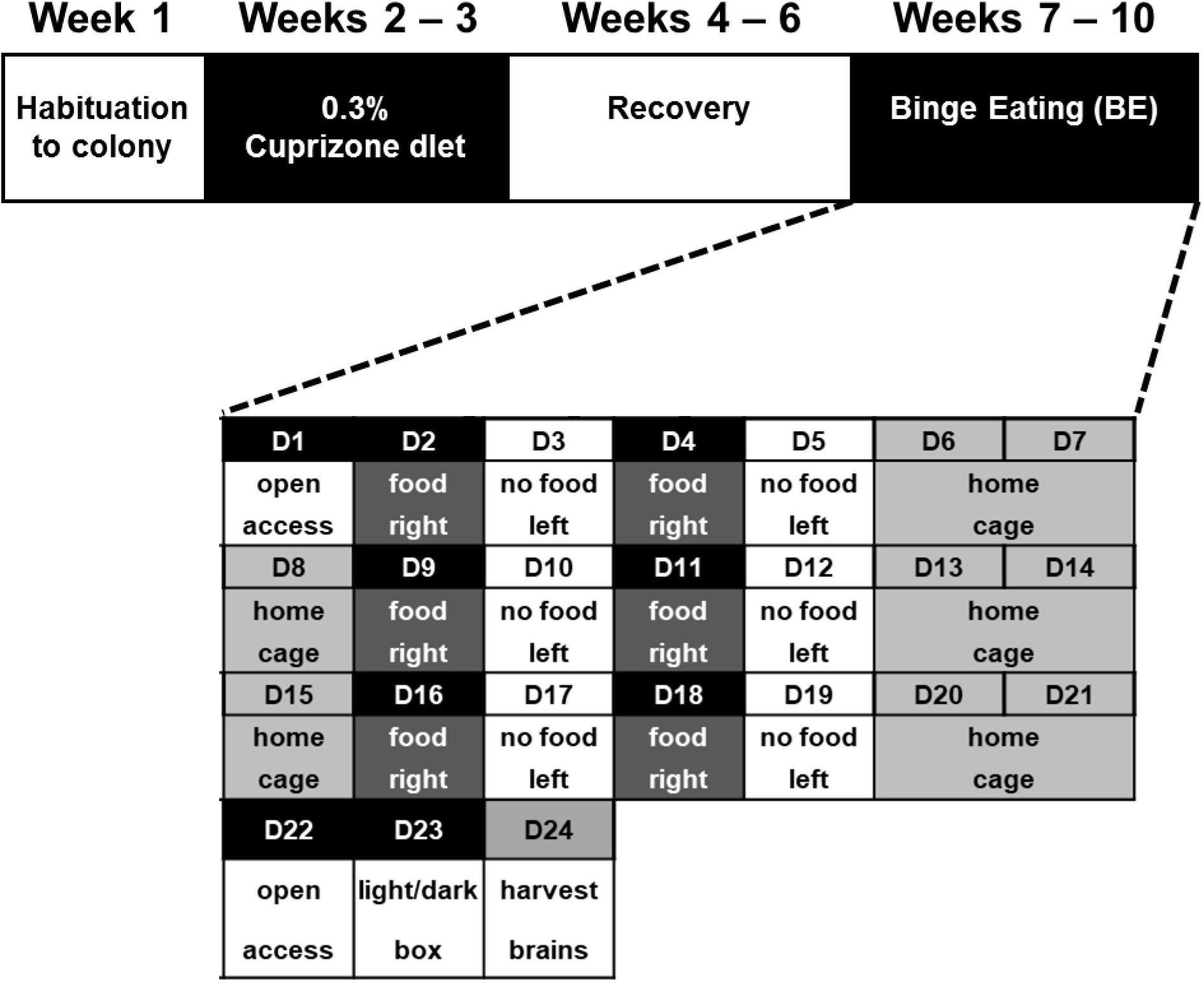
Study protocol and timeline. Mice were habituated for one week after their arrival. Mice then received either *ad libitum* standard laboratory chow or *ad libitum* cuprizone-treated lab diet for two weeks. After removal of cuprizone treatment, all mice were given *ad libitum* standard laboratory chow for three weeks. Finally, in weeks 7-10, training for binge eating (**BE**) commenced with continued *ad libitum* access to normal chow minus the 30 min training sessions with palatable food (**PF**). Mice were tested for side preference in the CPP chamber before and after training (D1 and D22) for BE of PF. On D2, 4, 9, 11, 16, and 18, mice were confined to the right side of the chamber with access to PF. On D3, 5, 10, 12, 17, and 19, mice were confined to the left side of the chamber with an empty food bowl. On D23, the light/dark conflict test was conducted for compulsive-like PF consumption. On D24, brains were harvested, flash frozen, and stored at − 80°C.

**Figure 2.**
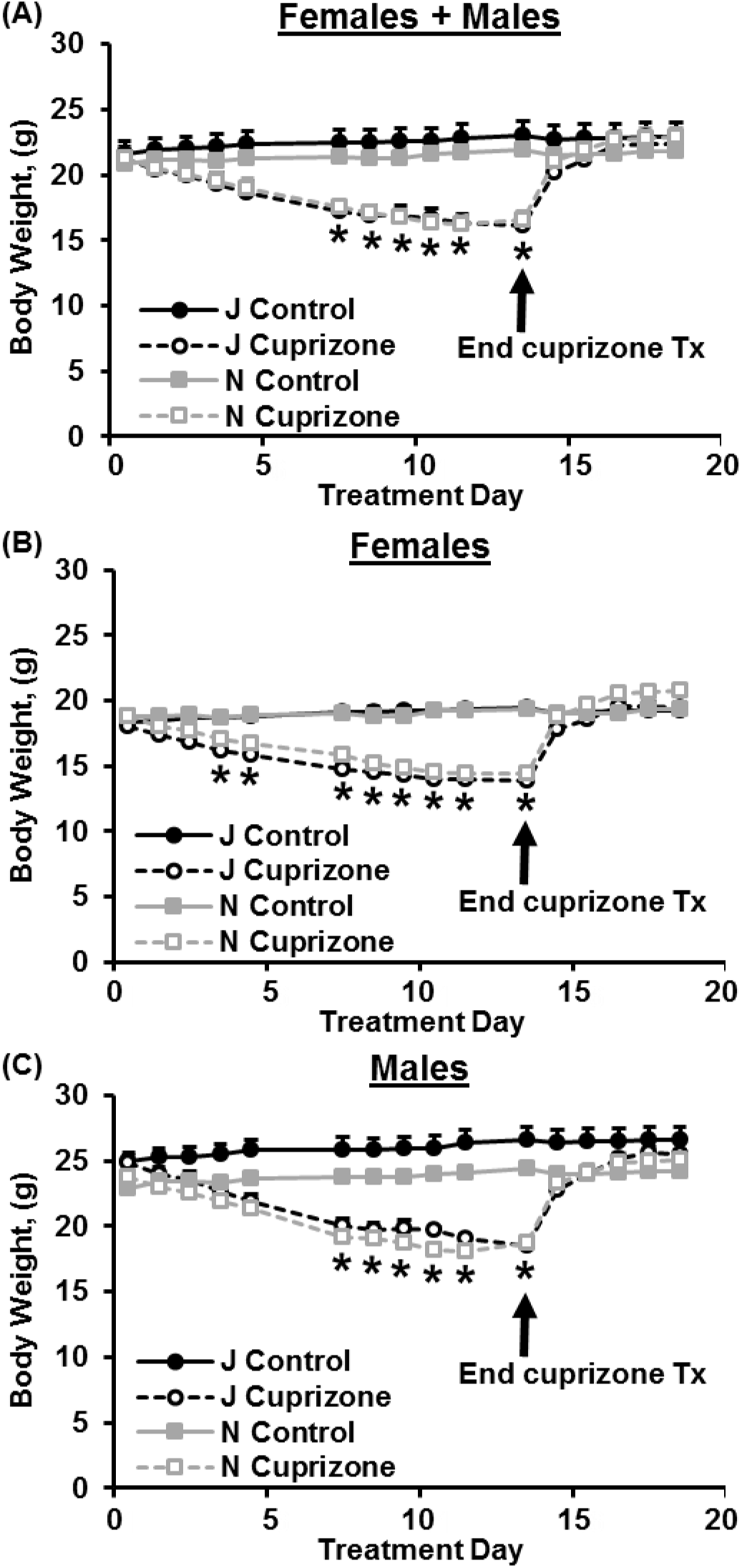
Changes in body weight over the course of cuprizone treatment and the week following termination of cuprizone treatment. **A)** In examining change in body weight, mixed model ANOVA with Substrain, Sex, and Treatment as factors and Day as a repeated measure indicated no significant effect of Substrain [F(1,55) = 1.68; p = 0.20]. However, there was an effect of Treatment [F(1,55) = 54.04; p = 1.0 × 10^−9^], Sex [F1,55) = 245.41; p < × 10^−16^] and a Substrain × Sex interaction [F(1,55) = 7.07; p = 0.01). There was also a main effect of Day [F(15,825) = 342.62; p < 2 × 10^−16^], a Substrain × Day interaction [F(15,825) = 2.02; p = 0.012], a Treatment × Day interaction [F(15,825) = 365.81; p < 10^−16^], a Sex × Day interaction [F(15,825) = 3.21; p =3.41 × 10^−5^], a Substrain × Treatment × Day interaction [F(15,825) = 2.73; p = 4.2 × 10^−4^], and a Treatment × Sex × Day interaction [F(15,825) = 7.24; p = 3.85 × 10^−15^]. Cuprizone treatment (D1-14) induced a decrease in body weight in both cuprizone-treated substrains relative to their respective control groups on D8-14 [unpaired t-tests: B6J cuprizone vs. B6J control: t(29) = 4.12, 4.44, 4.41, 4.65, 5.10, 5.50; *p = 2.8 × 10^−4^, 1.2 × 10^−4^, 1.3 × 10^−4^, 6.7 × 10^−5^, 1.9 × 10^−5^, 6.3 × 10^−6^; α_adjusted_ = 0.05/16 = 0.0031; B6NJ cuprizone vs. B6NJ control: t(30) = 4.52, 4.72, 5.10, 6.03, 5.95, 5.55; *p = 9.0 × 10^−5^, 5.1 × 10^−5^, 1.8 × 10^−5^, 1.6 × 10^−6^; α_adjusted_ = 0.0031]. **B)** In females, there was an effect of Treatment [F(1,28) = 47.21; p = 1.8 × 10^−7^] but no effect of Substrain [F(1,28) = 1.99; p = 0.17] and no interaction [F(1,28) = 1.85; p = 0.19]. Additionally, there was an effect of Day [F(15,420) = 226.0; p < 2.0 × 10^−16^], a nearly significant Substrain × Day interaction [F(15,420) = 1.68; p = 0.052], and a Treatment × Day interaction [F(15,420) = 239.03; p < 2 × 10^−16^]. Cuprizone-treated B6J females weighed less than control B6J females on D4-D14 [t(14) = 5.34, 5.79, 9.20, 9.23, 10.25, 12.08, 11.65, 12.42, **\***p = 1.1 × 10^−4^, 4.7 × 10^−5^, 2.6 × 10^−7^, 2.5 × 10^−7^, 6.9 × 10^−8^, 8.6 × 10^−9^, 1.4 × 10^−8^, 6.0 × 10^−9^,; α_adjusted_ = 0.0031]. Similarly, cuprizone-treated B6NJ females also weighed less than control B6NJ females on D4-D14 [t(14) = 4.46, 5.17, 6.53, 11.23, 11.45, 10.90, 10.64, 11.02; **\***p = 5.4 × 10^−4^, 1.4 × 10^−4^, 1.3 × 10^−5^, 2.2 × 10^−8^, 1.7 × 10^−8^, 3.2 × 10^−8^, 4.3 × 10^−8^, 2.8 × 10^−8^,; α_adjusted_ = 0.0031]. **C)** In males, there was a main effect of Substrain [F(1,488) = 48.5; p < 1.0 × 10^−11^], Treatment [F(1,488) = 170.3; p < 2 × 10^−16^], and Day [F(1,488) = 88.0; p = 0.0002] as well as a Substrain × Treatment interaction [F(1,488) = 12.4; p = 0.0005]. Cuprizone-treated B6J males weighed less than control B6J males on D8-D14 [t(13) = 5.29, 5.58, 5.41, 6.02, 6.63, 7.1; **\***p = 1.5 × 10^−4^, 9.0 × 10^−5^, 1.2 × 10^−4^, 4.3 × 10^−5^, 1.6 × 10^−5^, 8.1 × 10^−6^; α_adjusted_ = 0.0031]. Similarly, cuprizone-treated B6NJ males also weighed less than control B6NJ males on D8-D14 [t(14)= 6.55, 7.04, 7.60, 8.59, 7.48, 7.64; **\***p = 1.3 × 10^−5^, 5.9 × 10^−6^, 2.5 × 10^−6^, 5.9 × 10^−7^, 3.0 × 10^−6^, 2.3 × 10^−6^; α_adjusted_ = 0.0031].

When considering the female and male datasets separately, females showed significant weight loss earlier on at D4 and D5 (**Fig.2B**) and both sexes showed a significant reduction in body weight from D8-D14, after which there was recovery of weight loss within 24 following removal of the cuprizone diet and replacement with normal home cage chow (**Fig.2B,C**).

### Cuprizone treatment reduces BE and compulsive-like eating in male but not female mice of the BE-prone B6NJ substrain

Consistent with our previous report (Kirkpatrick et al., 2017), B6NJ mice showed greater overall PF consumption compared to B6J mice (**Fig.3**) - a behavior that we previously showed was associated with an enrichment of downregulated genes involved in myelination in the striatum (Kirkpatrick et al., 2017). However, contrary to our hypothesis that the demyelinating agent cuprizone would increase BE, prior treatment with cuprizone in the BE-prone B6NJ substrain actually decreased the amount of PF intake and decreased the slope of escalation in PF intake (**Fig.3A,B**). In contrast, there was no significant effect of cuprizone on the amount of PF intake in the BE-resistant B6J substrain, which, consistent with our previous findings (R. K. Babbs et al., 2018; Richard K. Babbs et al., 2019; Kirkpatrick et al., 2017), showed very little escalation of PF intake across days (**Fig.3A,B)**. Despite the lack of effect of cuprizone on the amount of PF intake in B6J mice, cuprizone induced a small but significant non-zero slope in escalation of PF intake across days (**Fig.3B**). In examining compulsive-like PF intake in the light/dark conflict test on D23, as expected, the BE-prone B6NJ strain showed a significant increase in PF intake relative to the BE-resistant B6J strain; however, there was no effect of cuprizone or interaction with substrain in the sex-combined dataset (**Fig.3C**).

**Figure 3.**
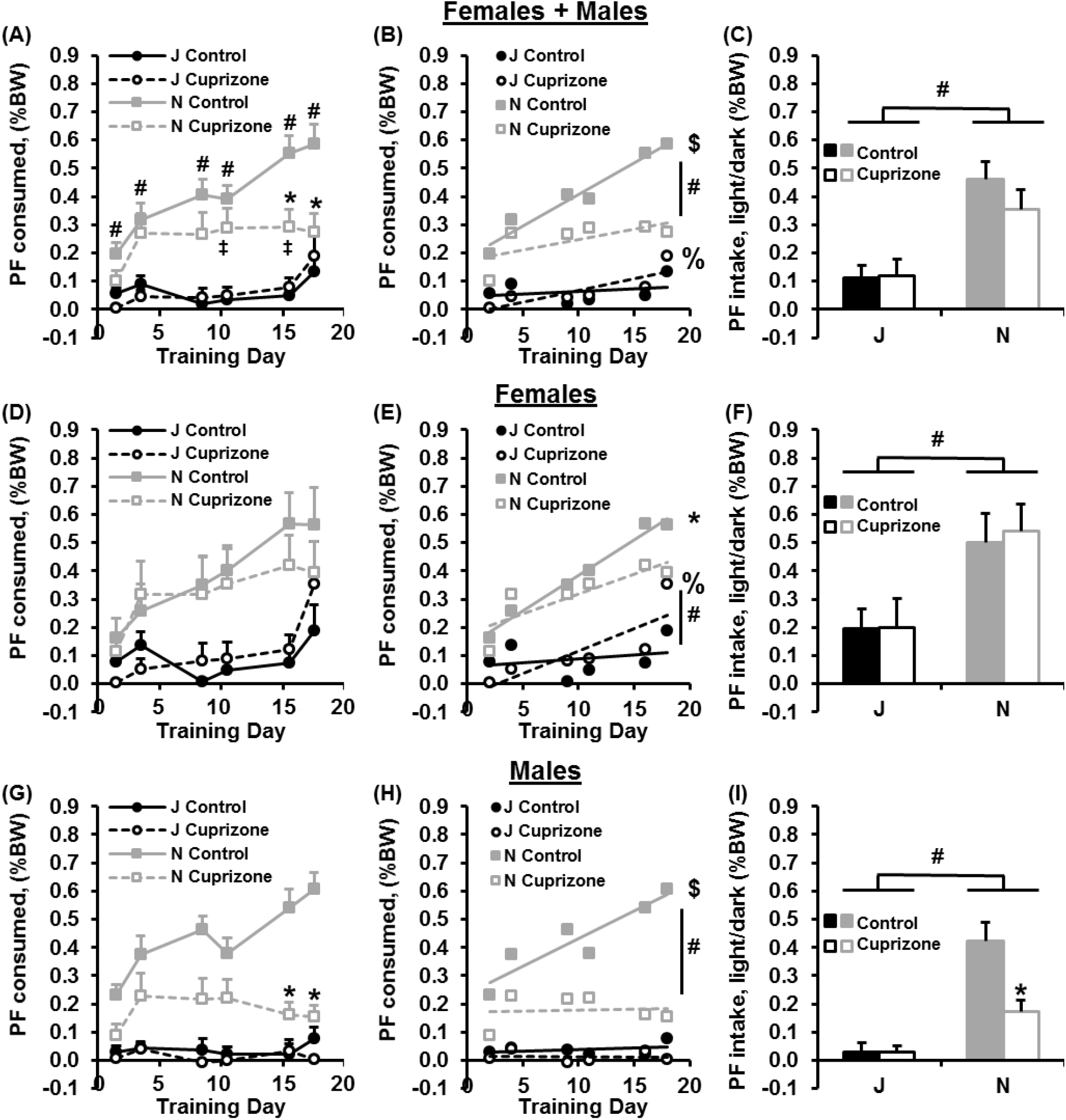
Cuprizone treatment reduces BE in male but not female B6NJ mice. **(A):** In examining the effect of cuprizone treatment on PF intake during BE training in the sex-combined dataset, a mixed effects ANOVA revealed a main effect of Substrain [F(1,55) = 50.16; p = 2.76 × 10^−9^], Treatment [F(1,55) = 4.56; p = 0.034], a Substrain × Treatment interaction [F(1,55) = 4.78; p = 0.033], and a nonsignificant effect of Sex [F(1,55) = 2.84; p = 0.098]. Additionally, there was an effect of Day [F(5,275) = 13.53; p = 8.2 × 10^−12^], a Substrain × Day interaction [F(5,275) = 4.83; p = 3.0 × 10^−4^], a Sex × Day interaction [F(5,275) = 2.89; p = 0.015], and a Substrain × Treatment × Day interaction [F(5,275) = 3.50; p = 4.37 × 10^−4^]. B6NJ control mice consumed more PF than both groups of B6J mice on all days (unpaired t-tests: **#**all ps < 0.003; α_corrected_ = 0.0083). Cuprizone-treated B6NJ mice consumed less PF than their control B6NJ counterparts on D16 and D18 [t(30) = 3.20, 3.49; *p = 0.0032, 0.0015, respectively; α_adjusted_ = 0.0083]. **(B):** In examining escalation in PF intake, both control B6NJ mice and cuprizone-treated B6J mice showed significant, non-zero slopes, indicating escalation [F(1,94) = 32.10, **$**p < 0.0001; F(1,88) = 7.15; **%**p < 0.01, respectively]. Furthermore, control B6NJ mice showed a significantly greater slope than cuprizone-treated B6NJ mice [F(1,188) = 6.07; **#**p = 0.015] but there was no difference in slope between treatments in B6J mice [F(1,182) = 2.83; p = 0.094]. **(C):** In examining compulsive-like PF intake in the light/dark conflict test, there was a main effect of Substrain [F(1,55) = 30.65; **#**p = 8.9 × 10^−7^ (B6NJ mice showed greater PF intake than B6J mice)] and Sex [F(1,55) = 14.07; p = 4.3 × 10^−4^], but no effect of Treatment [F(1,55) = 0.90; p = 0.35] and no interactions (ps > 0.17). **(D):** In examining the effect of cuprizone treatment on PF intake during BE training in females, there was a main effect of Substrain [F(1,28) = 14.63; p = 0.00067], but no effect of Treatment [F(1,28) = 0.081; p = 0.78] and no Substrain × Treatment interaction [F(1,28) = 0.51; p =0.48]. There was also an effect of Day [F(5,140) = 8.93; p = 2.2 × 10^−7^] and a Substrain × Day interaction [F(5,140) = 2.50; p = 0.033]. Importantly, the lack of effect of Treatment or interaction with Substrain was confirmed by a lack of significant pairwise difference in PF intake between cuprizone-treated B6NJ females and control B6NJ females for any of the six training days [t(14) = 0.55, 0.38, 0.19, 0.32, 0.96, 0.97; p = 0.59, 0.71, 0.85, 0.75, 0.35, 0.35, respectively; α_adjusted_ = 0.0083]. Thus, the reduction in PF intake in cuprizone-treated B6NJ mice in panels A-C is not mediated by females. **(E):**In examining escalation of PF intake in females, both the B6NJ control females and the cuprizone-treated B6J females showed a significant non-zero escalation [F(1,46) = 16.18, **\***p < 0.001; F(1,46) = 8.59, **%**p < 0.01, respectively] and although there was no difference in slope between the two treatments in B6NJ females [F(1,92) = 1.15; p = 0.29], cuprizone-treated B6J females showed a significantly greater slope than control B6J females [F(1,92) = 4.14; **#**p = 0.045] that was driven by the uptick in PF intake on the final training day (D16) in cuprizone-treated B6J females (D16). **(F):** In examining compulsive-like eating in the light/dark test in females, there was a main effect of Substrain [F(1,28) = 11.65; **#**p = 0.002] but no effect of Treatment [F(1,28) = 0.049; p = 0.83] and no interaction [F(1,28) = 0.031; p = 0.86]. The effect of Substrain was explained by greater overall PF intake in B6NJ females versus B6J females (**#**). **(G):** In examining the effect of cuprizone on PF intake during BE training in males, there was a main effect of Substrain [F(1,27) = 66.54; p = 9.2 × 10^−9^], Treatment [F(1,27) = 17.83; p = 2.5 × 10^−4^], and a Substrain × Treatment interaction [F(1,27) = 11.10; p = 0.0025]. There was also an effect of Day [F(5,135) = 5.85; p = 6.4 × 10^−5^], a Substrain × Day interaction [F(5,135) = 4.29; p = 0.0012], a Treatment × Day interaction [F(5,135) = 4.49; p = 8.06 × 10^−4^], and a Genotype × Treatment × Day interaction [F(5,135) = 2.99; p = 0.014]. Cuprizone-treated B6NJ males showed less PF intake than control B6NJ males on D16 and D18 [t(14) = 4.68, 6.31; p = 3.6 × 10^−4^, 1.9 × 10^−5^; α_adjusted_ = 0.0083], thus accounting for the decreased PF intake in the sex-collapsed dataset in panels A and B. **(H):** In examining the slopes in escalation, only the B6NJ control males showed a significant, non-zero escalation in PF consumption [F(1,46) = 22.83; **$**p < 0.0001] and this slope was significantly greater than cuprizone-treated B6NJ males [F(1,92) = 10.23; **#**p = 0.0019]. **(I):** In examining compulsive-like PF intake in the light/dark test in males, there was a main effect of Substrain [F(1,27) = 35.81; **#**p = 2.2 × 10^−6^ (B6NJ males showed greater PF intake than B6J males)], Treatment [F(1,27) = 8.31; p = 0.0077], and a Substrain × Treatment interaction [F(1,27) = 7.55; p = 0.011]. The interaction was explained by cuprizone-treated B6NJ males showing reduced compulsive-like PF intake compared to control B6NJ males [t(14) = 3.18; **\***p = 0.0067].

In examining females only, there was no significant effect of the cuprizone diet on PF consumption or compulsive-like PF intake (**Fig.3D,F**). However, there was a small but significant increase in the slope of escalation of PF intake in cuprizone-treated B6J females due to the uptick in PF intake on the final day of BE training (D18; **Fig.3D,E**). Furthermore, the escalation in cuprizone-treated B6J females was significantly greater than control B6J females (**#**p = 0.045; **Fig.3D**).

In examining males only, cuprizone-treated B6NJ mice showed a robust reduction in PF intake relative to their control B6NJ counterparts (**Fig.3G**). Additionally, cuprizone treatment eliminated the slope in escalation of PF intake (**Fig.3H**) and significantly reduced compulsive-like PF intake in B6NJ males in the light/dark conflict test (**Fig.3I**). Thus, the cuprizone-induced reduction of PF intake in the sex-combined dataset was completely accounted for by male B6NJ mice.

In examining locomotor activity on D23 in the light side of the light/dark test for compulsive-like eating, there was a main effect of Genotype (F1,55 = 4.72; p = 0.034) and Sex (F1,55 = 3.97; p = 0.051) but no effect of Treatment [F(1,55) = 0.03; p = 0.86) and no interactions (ps > 0.54) (data not shown). In examining time spent in the light-paired side, there was also no effect of Genotype [F(1,55) = 0.20; p = 0.66), Treatment [F(1,55) = 0.25; p = 0.62], or Sex [F(1,55) = 3.55; p= 0.065] and no interactions (ps > 0.15) (data not shown).

### Effect of cuprizone on locomotor activity on D1 during initial preference assessment and on D22 during PF-CPP assessment

We did not observe any evidence for a potential confounding influence of cuprizone treatment on locomotor activity that could explain differences in PF intake. Specifically, in considering the sex-combined dataset for D1 locomotor activity during initial preference for the PF-paired side (5 weeks after the beginning of cuprizone treatment or i.e., three weeks after the termination of cuprizone treatment), there was no effect of Substrain or interaction with Treatment (Fig.4A,C,E). In considering the sex-combined dataset for D22 locomotor activity during assessment of PF-CPP, there was a significant Genotype × Treatment × Sex interaction that was explained by a small but significant decrease in locomotor activity in cuprizone-treated B6J females (Fig.4B,D,F) in the absence of any significant effect of PF intake (see Fig.3D).

**Figure 4.**
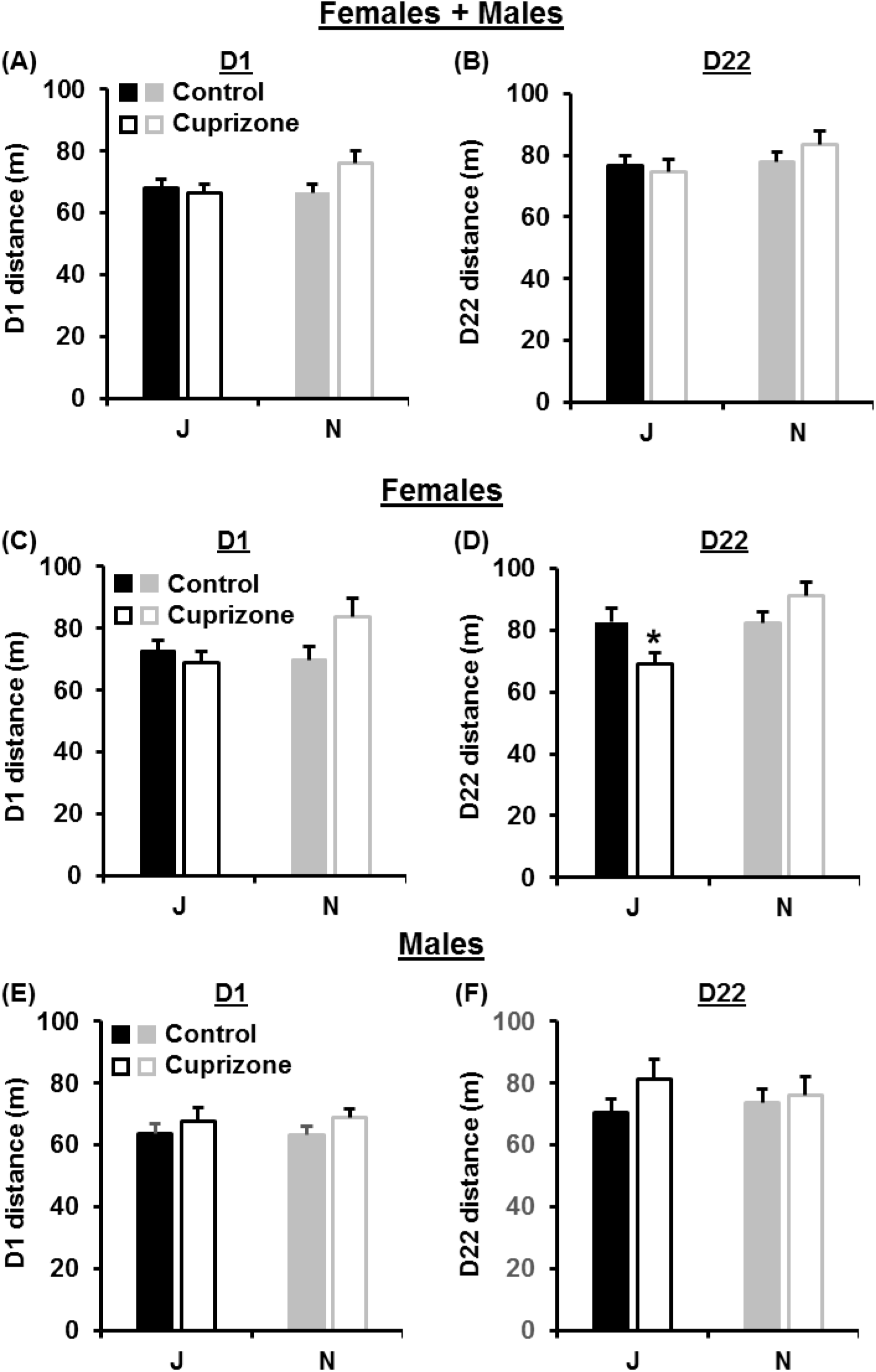
Effect of cuprizone treatment on locomotor activity. **(A,C,E):** In examining locomotor activity on D1 over 30 min, there was an effect of Sex with females showing the expected greater locomotor activity than males [F(1,55) = 7.84; p = 0.0071] but no effect of Genotype [F(1,55) = 1.29; p = 0.26], Treatment [F(1,55) = 3.08; p = 0.085], or any interactions (p > 0.086). **(B,D,F):** In examining locomotor activity on D22 over 30 min, there was no effect of Genotype [F(1,55) = 2.26; p = 0.14], Treatment [F(1,55) = 0.32; p = 0.57], or Sex [F(1,55) = 3.30; p = 0.075]. However, there was a significant Genotype × Treatment × Sex interaction [F(1,55) = 4.99; p = 0.030]. To determine the source of this three-way interaction on D22, we analyzed females and males separately. **(D):** For D22 locomotor activity in females, there was a main effect of Genotype [F(1,28) = 6.86; p = 0.014], no effect of Treatment [F(1,28) = 0.38; p = 0.55], and a significant Genotype × Treatment interaction [F(1,28) = 6.94; p = 0.014]. The source of the interaction was explained by cuprizone-treated B6J females showing less locomotor activity on D22 than control B6J females [t(14) = 2.23; p = 0.043]. For males, there was no effect of Genotype [F(1,27) = 0.016; p = 0.90], Treatment [F(1,27) = 1.46; p = 0.24], or interaction [F(1,27) = 0.61; p = 0.44].

In examining PF-CPP via the change in time spent on the PF-paired side between D1 and D22 (D22-D21; s), there was no effect of Genotype [F(1,55) = 0.01; p = 0.91], Treatment [F(1,55) = 0.47; p = 0.50], Sex [F1(55) = 0.96; p = 0.33], or any interactions (ps > 0.41). In an additional analysis of PF-CPP, we included Day as a repeated measure in a mixed effects ANOVA and in this case, we observed an overall effect of Day [F(1,55) = 6.74; p = 0.012] but no effect of Genotype [F(1,55) = 0.056; p = 0.82], Treatment [F(1,55) = 0.64; p = 0.43], Sex [F(1,55) = 0.69; p = 0.41], or any interactions (ps > 0.37) (data not shown).

### Effect of cuprizone on myelin proteins in the striatum

We previously reported a BE-induced downregulation of a gene network enriched for myelination – an effect that was driven by the BE-prone B6NJ substrain (Kirkpatrick et al., 2017). Here, we examined differences in the myelin proteins MAG and MBP in B6NJ versus B6J mice and the effect of cuprizone on these protein levels. There were no effect of cuprizone treatment or substrain on MAG levels (**Fig.5A,B**). However, for MBP, cuprizone treatment induced a significant decrease in total MBP and in each of the MBP isoforms compared to control mice (**Fig.5A,D**). There was also a main effect of substrain on MBP, with B6NJ showing a decrease in the 14 kDa MBP protein isoform (**Fig.5D**). Despite the main effects of Treatment and Substrain, there were no main effects of Sex or interactions with Sex (statistics are reported in the figure legend of **Fig.5**).

**Figure 5.**
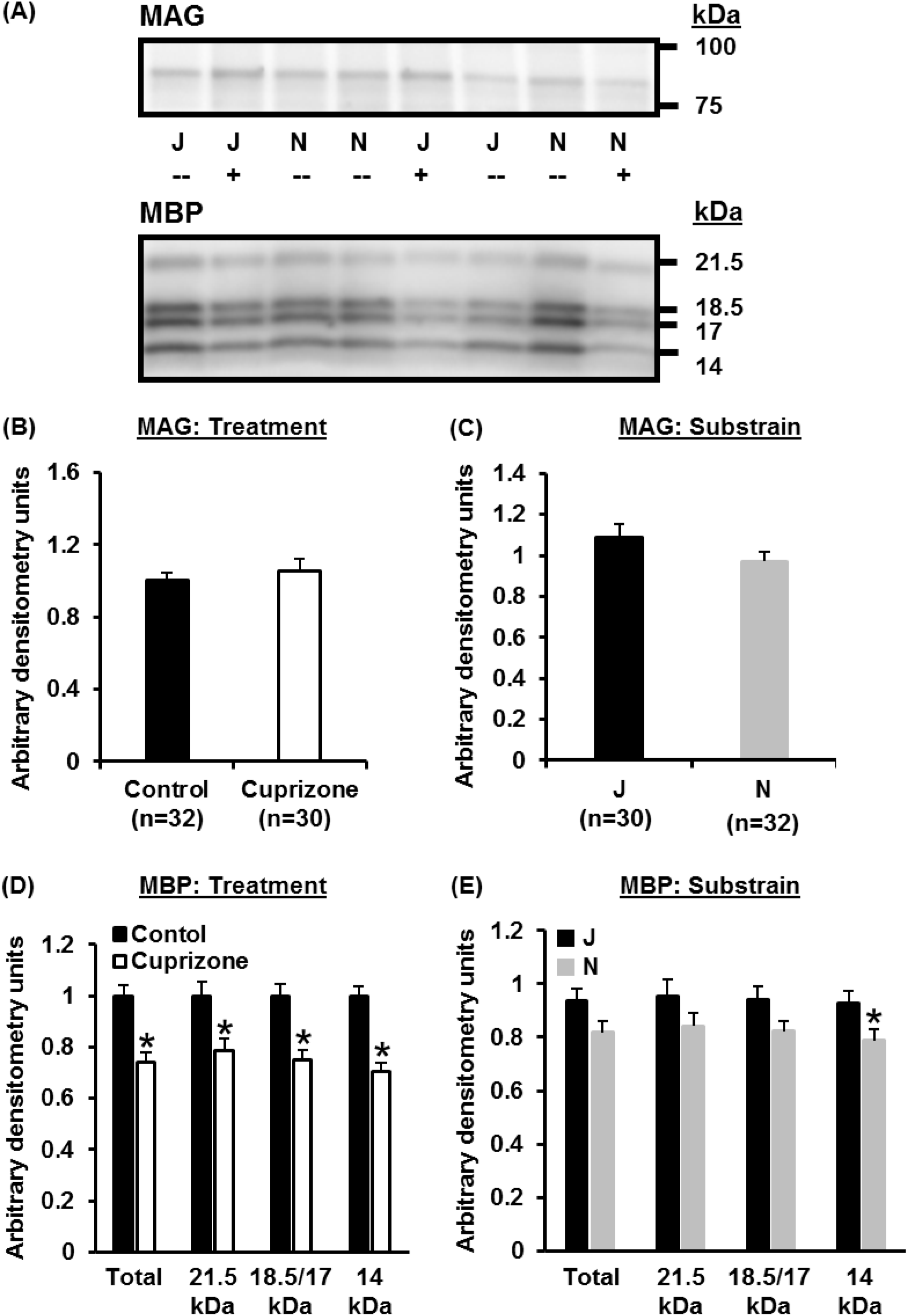
Effect of Substrain and Treatment on MAG and MBP protein levels in the striatum. **(A):** Representative immunoblots of striatal tissue from for MAG and MBP from cuprizone-treated (+) and untreated (-) C57BL/6J (**J**) and C57BL/6NJ (**N**) mice. **(B,C):** For MAG protein, there was no effect of Cuprizone Treatment [F(1,54) = 0.47; p = 0.49], Substrain [F(1,54) = 2.28; p = 0.14], Sex [F(1,54) = 2.87; p = 0.10] or any interactions (ps > 0.34). The lack of effect of Treatment and Substrain on MAG protein levels is shown graphically in panels B and C as a comparison with MBP (below). **(D):** <u>For total MBP protein</u>, there as an effect of Treatment [F(1,54) = 20.96; p = 2.8 × 10^−5^] and a nearly significant effect of Substrain [F(1,54) = 3.72; p = 0.059] but no effect of Sex [F(1,54) = 0.15; p = 0.70] and no interactions (ps > 0.14). <u>For the 21 kDa band</u>, there was a main effect of Treatment [F(1,54) = 8.06; p = 0.0064] but no effect of Substrain [F(1,54) = 1.90; p = 0.17], Sex [F(1,54) = 0.012; p = 0.91], or any interactions (ps > 0.10). <u>For the 18.5 + 17 kDa bands</u>, there was a main effect of Treatment [F(1,54) = 17.42; p = 0.00011] and a nonsignificant effect of Substrain [F(1,54) = 3.29; p = 0.075] but no effect of Sex [F(1,54) = 0.078; p = 0.78] and no interactions (ps > 0.15). <u>For the 14 kDa band</u>, there was an effect of Treatment [F(1,54) = 40.24; p = 4.8 × 10^−8^] and Substrain [F(1,54) = 7.8; p = 0.0072] but no effect of Sex [F(1,54) = 0.98; p = 0.33] and no interactions (ps > 0.11). **(D):** To follow up on the Treatment effects, unpaired t-tests for the total, 21 kDa, 18/17 kDa, and 14 kDa MBP bands indicated a significant decrease in MBP for all bands [t(60) = 4.56, 2.87, 4.19, and 6.00; p = 2.6 × 10^−5^, 0.0057, 9.3 × 10^−5^, 1.2 × 10 ^7^, respectively]. **(E):** To follow up on the trending Substrain effect, unpaired t-tests for the total, 21 kDa, 18.5/17 kDa, and 14 kDa MBP bands indicated a significant decrease in the immunostaining of the 14 kDa band (**\***) in the B6NJ strain (N) [t(60) = 1.82, 1.42, 1.76, and 2.32; p = 0.073, 0.16, 0.084, and **\***0.024, respectively].

## DISCUSSION

The demyelinating agent cuprizone induced comparable weight loss in females and males of both the BE-resistant B6J substrain and the BE-prone B6NJ substrain (**Figs.1,2**), yet it induced robust substrain- and sex-dependent effects on PF intake. In contrast to our hypothesis that administration of the demyelinating agent cuprizone would enhance BE, we observed the opposite result - a robust reduction in BE in male but not female mice of the BE-prone B6NJ strain (**Fig.3**). The male-selective decrease in BE in cuprizone-treated B6NJ males could not be explained by confounding effects on locomotor activity (**Fig.4**) or by sex-interactive effects in the level of MAG or MBP (**Fig.5**) – two major myelin proteins.

Dose-dependent, reversible body weight loss during dietary cuprizone administration is well-documented (Hiremath et al., 1998; Skripuletz, Gudi, Hackstette, & Stangel, 2011; Steelman, Thompson, & Li, 2012; Stidworthy, Genoud, Suter, Mantei, & Franklin, 2003). Severe weight loss contributes to the high mortality rates in cuprizone doses exceeding 0.3% (Hiremath et al., 1998; Stidworthy et al., 2003). The mechanism behind cuprizone-induced weight loss is unclear, but could involve copper chelation, given that copper deficiency in both rats (C. G. Taylor, Bettger, & Bray, 1988) and mice (Prohaska, 1983) results in reduced body weight. In the present study, we report a similar degree of weight loss in both female and male cuprizone-treated mice of both substrains (**Fig. 2**). Thus, it is unlikely that weight loss alone is responsible for the male-specific reduction in BE and compulsive-like eating in the BE-prone B6NJ substrain. Also, food restriction-induced weight loss promotes rather than reduces BE in rodents (Consoli, Contarino, Tabarin, & Drago, 2009; Pankevich, Teegarden, Hedin, Jensen, & Bale, 2010) and humans (Polivy, Zeitlin, Herman, & Beal, 1994), further suggesting that the reduction in PF consumption by cuprizone treatment is not likely caused by the weight loss that occurred three weeks prior to BE training.

In contrast to our original hypothesis, the demyelinating agent cuprizone produced very little effect in the BE-resistant B6J strain (although note the small but statistically significant increase in the slope of PF intake in cuprizone-treated B6J females versus their control B6J female counterparts; **Fig.3D**), nor did it enhance BE in the BE-prone strain (Kirkpatrick et al., 2017). Instead, cuprizone reduced BE in the BE-prone B6NJ strain and it did so only in male mice (**Fig.3**). This are several potential explanations underlying the substrain- and sex-dependent effect. Perhaps males are more sensitive to the ingestive and/or post-ingestive aversive properties of the cuprizone diet (e.g., taste, nausea) and show a learned, generalization effect on PF intake during BE. However, this explanation cannot account for the similar rate of body weight recovery in females and males following replacement of the home cage cuprizone diet with home cage chow (**Fig.2B,C**). Another possibility is that cuprizone induced a selective change in the perception or the hedonics of sweet taste in males. Indeed, only the cuprizone-treated B6NJ males initially consumed less PF than their control B6NJ male counterparts on the very first PF training day [t(14) = 2.47; p = 0.027; **Fig.3G**]. However, despite this initial reduction in PF intake, the cuprizone-treated B6NJ males showed a sharp escalation in PF intake from the first to the second training day, after which they plateaued rather than continuing to escalate like their control B6NJ male counterparts (**Fig.3G,H**). It is also possible that cuprizone induced an anhedonic-like state selectively in B6NJ males that caused a reduction in PF intake (Sen et al., 2019) or, e.g., novelty-induced hypophagia due to a depressive-like effect (Dulawa & Hen, 2005).

Here, we showed that two weeks of exposure to the cuprizone diet induced a significant reduction in the level of MBP when assessed at nearly nine weeks after the beginning of cuprizone without affecting MAG at the time of assessment, irrespective of Sex or Substrain (**Fig.5**). MBP is a major structural protein produced by oligodendrocytes and Schwan cells and redistributes to the processes to initiate axon ensheathment, compaction, thickening, and adhesion (Han, Myllykoski, Ruskamo, Wang, & Kursula, 2013; Harauz, Ladizhansky, & Boggs, 2009). Alternative splicing of MBP produces four major products coded by a single gene with alternative start sites for the larger and smaller isoforms, including the full length 21.5 kDa protein as well as the major 18.5 kDa isoform and the 17 and 14 Kd isoforms (de Ferra et al., 1985; Harauz et al., 2009). MAG is a 100 kDa transmembrane glycoprotein of the inner layer of the myelin sheath that is expressed in oligodendrocytes and Schwan cells and contributes to the formation and maintenance of the myelin sheath as well as inhibitory signaling cascades during neuronal regeneration (Han et al., 2013). Previous studies have found a decrease in both proteins following cuprizone treatment followed by a recovery of MAG and MBP levels during remyelination (Ludwin & Sternberger, 1984). It is unclear why we did not observe a change in MAG levels but perhaps the abbreviated protocol in our study was insufficient to change MAG levels. In support, a previous study of MBP and MAG in a viral model of recurring demyelinating lesions found a preferential decrease in MBP (Dal Canto & Barbano, 1985). Alternatively, our cuprizone regimen could have induced a reduction in MAG early on that recovered by the time of assessment at nearly nine weeks following the beginning of cuprizone treatment as the process of remyelination ensued (Ludwin & Sternberger, 1984). It is also possible that there are brain region-specific changes in myelin-associated proteins and that the striatum shows a different profile than, e.g., the corpus callosum which is the hallmark tissue that is typically employed to assess the degree of cuprizone-induced demyelination.

We observed a decrease in MBP in the BE-prone B6NJ substrain versus BE-resistant B6J substrain that was significant for the14 kDa isoform (**Fig.5**) and was consistent with our previous observation of an association of BE with the downregulated expression of a set of genes enriched for myelination (Kirkpatrick et al., 2017). In that study, the enrichment was identified as a function of Treatment (PF vs. Chow) and not as a function of Genotype (+/+ versus +/-) at the cytoplasmic FMR1-interacting protein 2 (*Cyfip2*) gene - the likely major causal genetic factor explaining B6 substrain differences in BE (Kirkpatrick et al., 2017; Kumar et al., 2013). However, a limitation of the current study is that because all mice received PF for BE training, we cannot state whether the decrease in MBP at the protein level in the B6NJ substrain versus the B6J substrain is simply a function of Genotype (Substrain) or if it also depends on training in BE with PF. Accumulating studies suggest a potential role of CYFIP proteins in myelination/demyelination. CYFIP2 facilitates the adhesion of T cells in CD4+ cells from patients with multiple sclerosis – a disease with the hallmark feature of demyelination (Mayne et al., 2004). A recent study employing overexpression of *Cyfip1* (gene homolog of *Cyfip2*) in mice resulted in changes in gene expression enriched for myelination (Fricano-Kugler et al., 2019). Furthermore, a second recent study found that *Cyfip1* haploinsufficiency disrupted white matter, myelin sheath (axon diameter), and the intracellular distribution of MBP from the cell body to the processes of mature oligodendrocytes without changing axon number or diameter (Silva et al., 2019). We recently reported that *Cyfip1* haploinsufficiency disrupts BE in a complex manner that depends on the *Cyfip2* genotype inherited from the particular B6 substrain as well as sex, and parent-of-origin (Richard K. Babbs et al., 2019). Future immunoblotting and immunohistochemical analyses will be necessary to determine if there are pre-existing differences in myelin proteins between naïve B6 substrains and if so, whether these differences are caused by the native *Cyfip2* coding mutation in the B6NJ substrain (Kirkpatrick et al., 2017; Kumar et al., 2013).

An additional factor that could contribute to sex differences in the effect of cuprizone on BE is weight loss-induced rebound hyperphagia. We are unaware of any studies demonstrating sex differences in eating behavior after weight loss in rodents – with or without cuprizone. However, greater hyperphagia has been reported in females compared to males after hypothalamic lesions in rats (Valenstein, Cox, & Kakolewski, 1969) and *Il18* deletion in mice (Zorrilla et al., 2007). Moreover, when rats were acutely fasted (12 hours), males showed a modest increase in food consumption in the 24 h post-fast, while females showed a robust increase in consumption that was significantly greater than that of the males (Gayle, Desai, Casillas, Beloosesky, & Ross, 2006). In a closer examination of our data, although both females and males treated with cuprizone showed similar reductions in body weight and similar regaining of body weight to pre-cuprizone levels (**Fig.2**), B6NJ females showed a faster percent body weight increase compared to B6NJ males during the first 24 h after the cuprizone diet was replaced with the normal chow diet [KEITH: t(df)= 2.5; p = 0.02; data not shown), providing some evidence for slower rebound hyperphagia in B6NJ males that could extend to less acute and escalated PF intake in the 30 min training trials.

Another potential factor explaining the male-specific reduction in BE with cuprizone could involve sex differences in the demyelinating effects of cuprizone and/or the process of remyelination. Previous reports have found no significant sex difference in cuprizone-induced demyelination in C57BL/6J mice (L. C. Taylor, Gilmore, Ting, & Matsushima, 2010). However, a separate study in the SJL inbred mouse strain showed less severe cuprizone-induced demyelination and loss of oligodendrocytes in female SJL and Swiss Webster mice (Ludwin, 1978; L. C. Taylor, Gilmore, & Matsushima, 2009) which is consistent with the lack of significant behavioral effects observed with B6NJ females in the current study. Given that we used a different cuprizone regimen and importantly, given that we observed a sex difference in cuprizone behavioral effects in a different genetic B6 substrain (B6NJ), it is possible that B6NJ female mice are less sensitive to cuprizone-induced demyelination which in turn could explain the lack of behavioral effect. However, our current immunoblotting results with MAG and MBP fail to support this hypothesis as we did not identify any significant Sex effects or Sex-interactive effects at nearly nine weeks after the beginning of cuprizone treatment. Assessment of myelin proteins via immunoblot and immunohistochemical procedures at multiple earlier time points (e.g., starting at five weeks) (Doan et al., 2013) and in multiple brain regions could address whether the enhanced behavioral effects of cuprizone in B6NJ males are supported by enhanced or earlier demyelination.

To summarize, we observed a male-specific reduction in BE in the BE-prone B6NJ substrain in response to the demyelinating agent cuprizone. These results suggest that sex-specific neurobiological mechanisms could underlie BE in a manner that depends on genetic background. Future studies employing additional and more selective demyelination/remyelination strategies in a spatiotemporal manner on multiple genetic backgrounds are necessary to test the contribution of sex differences in myelin dynamics in the establishment of and recovery from BE.

## ACKNOWLEDGEMENTS

We thank Dr. Carmela R. Abraham for a discussion regarding the use of the cuprizone model in mice. This study was supported by R21DA038738, Spivack Award (Boston University), NIDA Summer Diversity Fellowship Program, and Boston University’s Transformative Training Program in Addiction Science (Burroughs-Wellcome #1011479).

## AUTHOR CONTRIBUTIONS

R.K.B. helped design the study and write the manuscript as well as treated the mice with cuprizone and analyzed behavior. J.A.B. oversaw and ran the immunoblots and conducted the immunoblot analysis and analyzed behavior. J.C.K. ran the behavioral study and contributed to the analysis. R.M., J.A., and A.S. ran immunoblots and contributed to the analysis. E.J.Y. performed protein extractions and protein quantification. M.M.C. conducted video tracking and behavioral analysis. C.D.B. designed and oversaw the study and wrote the manuscript.

